# The molecular basis of hard cornification in the keratinized jaw sheaths and keratodonts of anuran tadpoles

**DOI:** 10.64898/2026.09.30.755311

**Authors:** Pedro Henrique dos Santos Dias, Kevin Hsiung, Natalie Stallard, Jackson Jones, Andrew Capitosti, Sean Harrington, Paul Lukas, Ioannis Chrysostomakis, Florencia Vera Candioti, Oliver Hawlitschek, Alexander Haas, Astrid Böhne, Daniel J. Paluh

## Abstract

Hard cornification is the developmental process that produces the hardened epidermal structures that characterize tetrapod diversity, including scales, feathers, hair, and claws. This process relies on keratin proteins and additional reinforcing proteins that assemble, cross-link, and strengthen tissues to become mechanically resilient. Although the molecular composition of hard cornified tissues is known to vary among tetrapods, the program is least understood in amphibians. Within frogs, hard cornification occurs in the mouthparts of most tadpoles and in the nuptial pads, hindfoot claws, and foot tubercles of some adults. We characterized transcriptomic profiles of *Bombina orientalis* tadpoles at three developmental stages before, during, and after mouthpart cornification. We identified 25 keratin genes orthologous to amniote hard keratins in the *B. bombina* genome, 18 of which are upregulated in cornifying larval mouthparts. Furthermore, two of these hard keratin paralogs and an amphibian epidermal differentiation gene exhibit distinct spatial expression patterns within the jaw sheaths and keratodonts evidenced by *in situ* hybridization, suggesting they have discrete functional roles. The newly discovered diversity of the hard keratin repertoire in *Bombina*, and the concurrent expression of many of these paralogs in tadpole mouthparts, suggests that anuran hard cornification is much more complex than adult cornified traits thus far characterized in *Xenopus* frogs. Our results also challenge the notion that the diversification of keratins in Amphibia is primarily associated with the transition to more terrestrial habits, and may instead be shaped, in part, by the novel feeding apparatus of aquatic tadpoles.

## 1 Introduction

A hallmark of tetrapod evolution is the diversity of hardened epidermal structures, including hair, scales, feathers, and claws, that provide protection, insulation, and mechanical defence. Described as keratinized appendages, these structures are a product of the terminal stage of keratinocyte differentiation, known as hard cornification, in which living keratinocytes undergo programmed cell death and become mechanically resilient corneocytes (Bragulla and Homberger 2009; Alibardi 2022a). Hard cornification depends on keratin intermediate filament proteins (hereafter keratins) that assemble the keratin intermediate filament scaffold, as well as additional reinforcing proteins that cross-link and fortify the scaffold (Holthaus et al. 2025). Keratin-like genes arose during early metazoan evolution but rapidly diversified in sarcopterygians, corresponding to the water to land transition (Ho et al. 2022). Functional keratin intermediate filaments form through the polymerization of one type I (acidic) and one type II (basic-to-neutral) keratin protein into coiled-coil structures (Vandebergh and Bossuyt 2012; Lee et al. 2012). Although this keratin architecture is conserved, the additional reinforcing proteins that harden cornified tissues differ substantially among amphibians, mammals, and sauropsid reptiles (Alibardi et al. 2009; Alibardi 2010). Hard cornification is the least understood in Amphibia (Holthaus et al. 2025), which has limited our understanding of how hardened epidermal structures diversified across tetrapods.

In both mammals and sauropsids, a key regulatory component of hard cornification is the transcription factor Hoxc13, which activates specialized type I and type II hard keratin genes (i.e., hair keratins; Eckhart et al. 2008; Carron et al. 2024). These hard keratins encode cysteine-enriched proteins that build the scaffold of hard cornified structures, including hair and nails. This hard keratin gene subcluster has expanded and contracted across lineages, corresponding to the elaboration or reduction of cornified traits. For example, the hard keratin repertoire contracted in hairless cetaceans (Nery et al. 2014) and was entirely lost in clawless snakes (Ehrlich et al. 2020). Amniote hard cornification also depends on proteins encoded by the epidermal differentiation complex (EDC), a conserved genomic locus encoding the calcium-binding S100 proteins, S100 fused-type proteins (SFTPs) that cross-link keratin filaments, and simple epidermal differentiation complex proteins (SEDCs) that form the cornified envelope (Strasser et al. 2014). Although mammals and sauropsids inherited an ancestral EDC, each lineage subsequently evolved additional structural proteins that contribute to the mechanical properties of hardened epidermal tissues. Mammals evolved keratin-associated proteins (KAPs), a diverse family of short, low-complexity, proteins enriched in sulfur-containing cysteine residues (Wu et al. 2008; Khan et al. 2014). Disulfide cross-linking among KAPs and keratins likely reinforces the filament-matrix scaffold, producing the highly rigid properties of mammalian hardened appendages (Plowman et al. 2021). KAPs likely originated through duplication and divergence of ancestral keratin genes (Eckhart and Ehrlich 2018), and their expression is also regulated by Hoxc13 (Pruett et al. 2004). In contrast, sauropsid reptiles evolved corneous beta-proteins (CBPs), a family of structural proteins that coat and reinforce the keratin scaffold during cornification (Holthaus et al. 2019). CBPs derive their mechanical strength from a conserved β-sheet and evolved through duplication and divergence of ancestral EDC genes (Calvaresi et al. 2016; Holthaus et al. 2019). Hard cornification cross-linking can occur through disulfide bonds catalyzed by Sulfhydryl oxidase (SOXase) enzymes and isopeptide bonds catalyzed by transglutaminase enzymes (Alibardi 2022b).

Amphibians generally possess soft, highly permeable, glandular skin specialized for cutaneous respiration and water uptake (Larsen 2021). As a result, hard cornification has received comparatively little attention in this clade, despite the evolution of numerous specialized hardened epidermal structures in frogs and salamanders (batrachians; Figure 1). The third living amphibian order, the caecilians, entirely lacks hard cornified structures (Torres-Sánchez et al. 2020). Like all vertebrates, amphibians possess type I and type II keratin gene families that are broadly expressed in epithelial tissues (Vandebergh and Bossuyt 2012; Suzuki et al. 2017). Frogs also possess a key regulatory component of amniote hard cornification: two keratin genes orthologous to amniote hard keratins, *krt34* (type I) and *krt59* (type II), are regulated by Hoxc13 and are expressed in cornifying claws and nuptial pads in *Xenopus laevis* and *X. tropicalis* (Carron et al. 2024; Steinbinder et al. 2026). Unlike their amniote counterparts, however, *Xenopus krt34* and *krt59* are not enriched in cysteine residues. These amphibian hard keratins have undergone lineage-specific expansion in at least one species with elaborated cornified appendages—the Leishan moustache toad (*Leptobrachium leishanense*)—which has been linked to the prominent keratinized spines along the upper jaw in males (Li et al. 2019).

**FIGURE 1.**
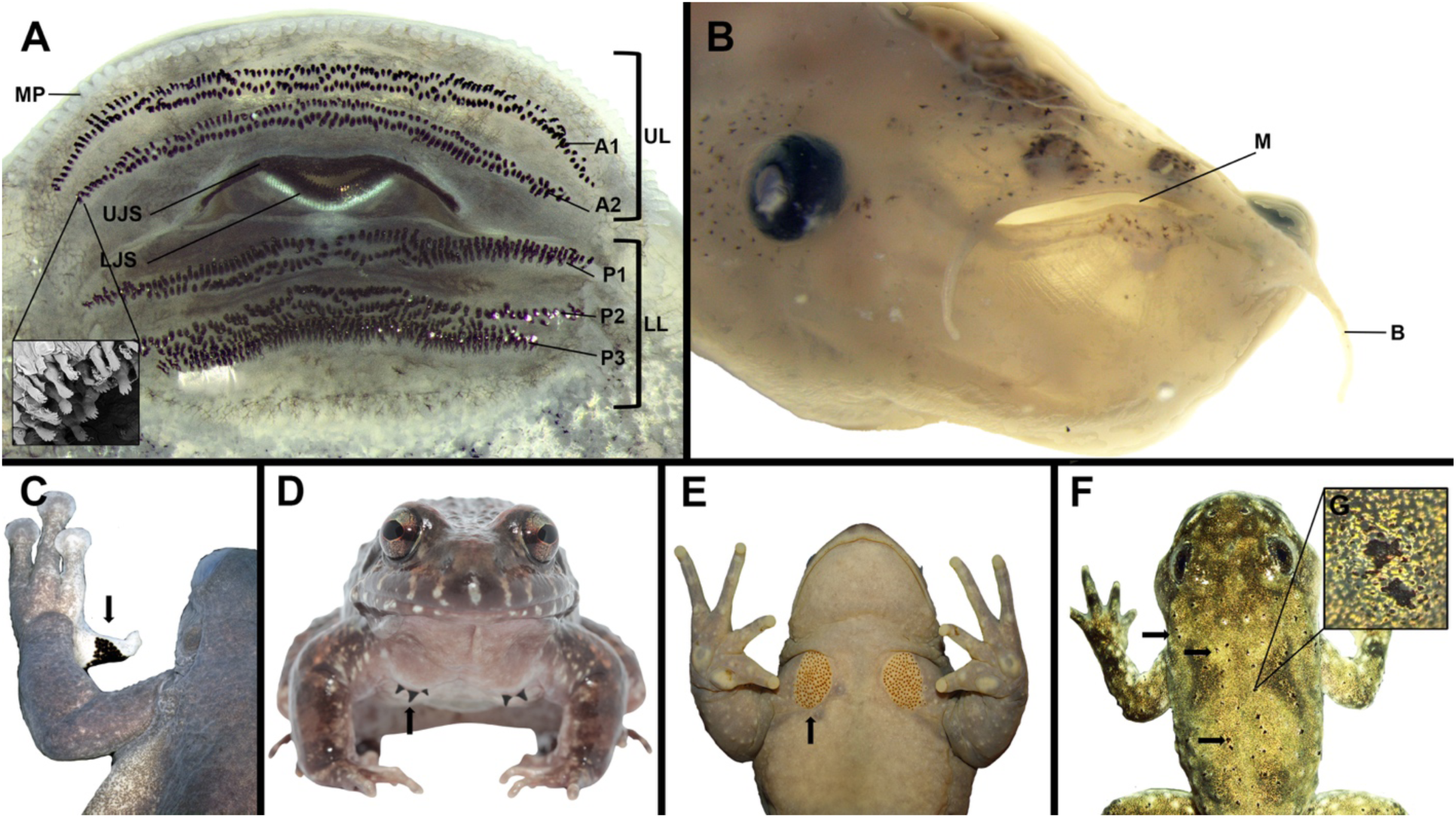
(A) Oral disc of *Bombina variegata* tadpole composed of keratinized jaw sheaths and keratodonts. (B) Simple mouth opening of *Xenopus laevis* tadpole lacking cornified mouthparts. Examples of cornified traits in postmetamorphic frogs include nuptial pads on the forelimbs (C, adult male of *Atlanthyla spinipollex* UF 166704) and chest (D, adult male of *Leptodactylus syphax*; E, adult male of *Alsodes gargola*, MACN 38891) and body tubercles (F, *Bombina variegata* froglet). Courtesy photos from Paulo Durães Pinheiro (C), Tiago L. Pezzuti (D), and Boris L. Blotto (E).

The additional reinforcing proteins that harden cornified tissues in amphibians have remained poorly understood. Like other vertebrates, amphibians possess the ancient core of the EDC, including calcium-binding S100 proteins (Zimmer et al. 2012). Previously, additional EDC-encoded proteins associated with structural reinforcement, including SFTPs and SEDCs, were thought to represent amniote innovations. However, two SFTP genes are present in the genomes of *Xenopus*, a salamander (*Ambystoma mexicanum*, the axolotl), and several caecilians (Mlitz et al. 2017; Sachslehner and Eckhart 2022). However, it remains unknown whether these proteins are incorporated into hard cornified tissues or contribute to structural reinforcement. Caecilians are the only amphibians known to possess SEDC genes (Sachslehner and Eckhart 2022), yet the hard cornification program likely decayed in this lineage through the loss of *krt34* and *krt59* (Carron et al. 2024). A newly discovered gene family, the amphibian epidermal differentiation cluster (AEDC), likely plays a key role in the structural reinforcement of hard cornified tissues in frogs (Sachslehner et al. 2026). A subset of these AEDCs encode short, cysteine-enriched, low-complexity proteins that may be functionally analogous to mammalian KAPs in enabling disulfide cross-linking. For example, *aedc13* is spatially co-expressed with *krt59* within epidermal nuptial pad cells of *Xenopus* undergoing cornification (Sachslehner et al. 2026; Steinbinder et al. 2026).

Amphibian hard cornification is therefore distinct from amniotes in several respects and requires further characterization across both lineages and trait diversity. In frogs, cornified structures can be found both in larvae and adults, including the mouthparts of tadpoles. The typical tadpole feeding apparatus, known as the oral disc, is composed of keratinized jaw sheaths (i.e., horny beak), several rows of keratinized labial teeth (i.e., keratodonts) above and below the jaw sheaths, and marginal papillae on an upper and lower labium (Figure 1; Altig 2007a; Dreger et al. 2025). In adults, nuptial pads, body tubercles, hindfoot claws, and foot tubercles can undergo hard cornification (Figure 1; Luna et al. 2018, Holthaus et al. 2025). Some of these traits, such as larval mouthparts and adult nuptial pads, are widely distributed across frogs (Luna et al. 2018; Vera Candioti et al. 2025), with significant phenotypic and ecological variation (Altig and Johnston 1989; Luna et al. 2018; Dias and Anganoy-Criollo 2024), whereas other cornified structures are restricted to particular clades. Most work on the molecular basis of amphibian cornification to date has been conducted on postmetamorphic structures (e.g., nails and nuptial pads in *Xenopus*, jaw spines in *Leptobrachium*), whereas little is known about the developmental genetics of tadpole mouthparts (Paluh et al. 2025).

The jaw sheaths of anuran larvae are composed of epithelial columnar basal keratinocytes that differentiate into stratified, flattened sheath cells that form the lingual and labial surfaces of the keratinized sheath, as well as central, stacked columns of cone-shaped cells that form the serrated, cutting edge of the sheath (Luckenbill 1965; Kaung 1975). Highly proliferative pre-cone cells differentiate into cone cells, and each cone cell column forms a single micro-serration along the jaw sheath edge. The keratodonts of tadpoles are similarly arranged as a stacked column of keratodont cells, and they continuously proliferate, cornify, and replace the apicalmost, worn structures throughout the life of a tadpole due to a highly regenerative mitotic zone of basal keratinocytes (Luckenbill 1964, 1965; Kaung 1975; Alibardi 2025). Each keratodont locus can therefore be composed of multiple erupted keratodont generations (Altig 2007a), although only the apicalmost cell comes into contact with feeding surfaces. During the process of hard cornification, the apicalmost jaw sheath and keratodont cells become black or dark brown in coloration.

In this study, we characterized the keratin repertoire of the *Bombina bombina* genome using molecular phylogenetics and generated transcriptomic profiles of *B. orientalis* tadpoles at three developmental time points to identify genes associated with hard cornification of larval mouthparts. We further assessed the spatial expression of two keratin genes and one AEDC gene using *in situ* hybridization in *Bombina variegata*, a close relative to *B. orientalis.* We found that *Bombina* has many keratin copies orthologous to the hard keratins of amniotes expressed in cornifying larval mouthparts, unlike the single *krt34* and *krt59* expressed in the claws of *Xenopus*. Furthermore, two of these hard keratins and an AEDC exhibited distinct spatial expression patterns within the jaw sheaths and keratodonts, suggestive of discrete functional roles during mouthpart cornification.

## 2 Material and Methods

### 2.1 Identification and characterization of keratins in the *Bombina bombina* genome

Because many keratin genes were uncharacterized in the *Bombina bombina* reference genome annotation, we used a Hidden Markov Model (HMM) search in HMMER (v. 3.4; http://hmmer.org/) to query for these loci. We downloaded known type I and II keratin sequences from Uniprot for human and mouse. For each of type I and type II keratins, we combined human and mouse sequences into a single fasta file, then aligned sequences with MAFFT v. 7.526; Katoh and Standley 2013) and trimmed poorly aligned regions with Trimal (v. 1.5.rev1; Capella-Gutiérrez et al. 2009). We then built an HMM for each keratin cluster (type I and type II) and used the HMM to search the *B. bombina* genome assembly (NCBI RefSeq assembly GCF_027579735.1), as well as the mouse genome assembly (NCBI RefSeq assembly GCF_000001635.27) to validate that we found expected hits. Because all keratins are structurally similar, searches for type I keratins hit many type II keratins, and *vice versa*. Therefore, we used a custom python script to classify proteins that were identified by both HMM searches based on which HMM provided a better hit. We used the R package gggenes (v. 0.7.0; Wilkins 2025) to visualize the spatial organization of keratin genes in *B. bombina*. Lastly, we used the R package Biostrings (v. 2.76.0; Pagès et al. 2025) to calculate cysteine richness for each keratin protein.

### 2.2 Molecular phylogenetics of keratins

Predicted proteins for every type I and type II keratin identified in the *Bombina bombina* genome, as well as the keratins previously identified in *Leptobrachium leishanense* genome (Li et al. 2019) were added to the vertebrate keratin type I and type II intermediate filament rod domain alignments of Carron et al. (2024) using the --keeplength option in MAFFT (v. 7.526). Our alignments therefore contain keratin sequences from a lungfish (*Protopterus annectens*), a caecilian (*Rhinatrema bivittatum*), a salamander (*Ambystoma mexicanum*), the western clawed frog (*Xenopus tropicalis*), *B. bombina*, *L. leishanense* and *Homo sapiens*. Maximum-likelihood phylogenies were inferred separately for the type I and type II keratin datasets in IQ-TREE (v. 3.1.1; Wong et al. 2026). ModelFinder was used to select the best-fitting amino acid substitution model according to the Bayesian information criterion (BIC), identifying LG+I+G4 as the optimal model for both datasets. Node support was assessed using 1,000 ultrafast bootstrap (UFBoot) replicates with bootstrap NNI optimization.

### 2.3 Specimen sampling for RNA-sequencing

We characterized the transcriptomic profiles of the developing mouth in *Bombina orientalis*. All *B. orientalis* embryos and tadpoles obtained for this study were produced at the breeding facility maintained at the Zoology Department at the Friedrich-Schiller-University Jena, Germany (animal welfare protocol of Friedrich-Schiller-University Jena). Embryos were staged according to the simplified staging table for anuran embryos and larvae (Gosner 1960). Three embryos were sampled at three different developmental stages: 1) during primary mouth formation (Gosner stages 19–20; stomodeum phase), which corresponds to the invagination of the primordial mouth followed by the rupture of the oropharyngeal membrane; 2) during the development of the oral apparatus (Gosner stages 21–22), characterized by the differentiation of the labia, jaw sheaths, and labial ridges, on which keratodonts develop; and 3) when the oral apparatus is fully formed (Gosner stages 23–24), presenting all the typical structures of the oral disc, including cornified jaw sheaths and keratodonts (Figure S1). Embryos and tadpoles were euthanized using buffered 1% tricaine methane sulfonate (MS-222) and individually placed in a sterile Petri dish with RNAlater Stabilization Solution (Thermo Fisher Scientific). The entire oral region was manually isolated using micro tweezers and stored at -80°C.

### 2.4 RNA extraction and sequencing

Total RNA was extracted from embryos using the RNeasy Mini Kit (Qiagen) following the manufacturer instructions. Subsequently, strand-specific library preparation and sequencing was performed by BMK Genomics, Münster, Germany. Sequencing data have been deposited in the INSDC under NCBI BioProject Number (pending).

### 2.5 Sequencing reads quality filtering, mapping and read counting

Raw reads were trimmed of adapters and quality filtered with Trimmomatic (v. 0.39; Bolger et al. 2014) with a sliding window size of 4-bp, a required minimum window quality of 15, and a minimum read length of 100 bp (⅔ of average read length). The quality of read data was assessed before and after trimming using FastQC (Andrews, 2010) and MultiQC (Ewels et al. 2016). A chromosome-scale *Bombina bombina* reference genome assembly (RefSeq assembly version GCF_027579735.1; BioProject: PRJNA915412) was indexed using STAR (v. 2.7.10a; Dobin et al. 2013) with the parameter “--sjdbOverhang 149” (read length - 1) as recommended by STAR. Trimmed reads were subsequently mapped to the indexed genome with STAR using the parameters “--quantMode GeneCounts --twopassMode Basic outFilterMultimapNmax 1” to produce both alignments in BAM format and count data. Under these parameters, STAR applies splice junction information generated in an initial mapping step to a second mapping run to reduce false positive junctions. Strandedness was considered when producing gene level counts. Mapping statistics were taken directly from the STAR output logs. Total number of input reads and mapped reads are detailed in Table S1.

### 2.6 Differential gene expression analysis

Count data was processed and analyzed with R v4.0.3 (R Core Team, 2013) using the DESeq2 package (v. 1.30.1; Love et al. 2014). Following the recommendations of DESeq2, lowly expressed genes were filtered out, retaining genes with a minimum read count of five for at least three samples. After this removal, we obtained a total dataset of 16,219 expressed genes. Principal component analysis (PCA) was performed for all replicates on the variance stabilizing transformation (VST) normalized expression data as implemented in DESeq2. Visualizing the PCA, we identified one replicate (BO_MO_C_2_26) that clustered inconsistently with the expected stage of oral disc development. Hierarchical clustering of the top 500 most expressed genes using the R package pheatmap (v. 1.0.12; Kolde and Kolde 2015) likewise showed replicates grouping according to stage of formation except for BO_MO_C_2_26. To avoid potential bias from a mislabeled or contaminated sample, this sample was excluded from all further analysis.

To determine significant differences in gene expression between developmental stages, count data was normalized using DESeq2’s internal median of means method, and normalized counts were fitted through generalized linear models (GLM) with a negative binomial distribution. *P*-values were then calculated through a Wald-test and corrected for multiple testing with the Benjamini and Hochberg method in DESeq2. We retrieved log2 fold changes in expression and their corresponding *P*-value for all possible pairwise comparisons (stomodeum vs. during oral disc formation; during oral disc formation vs. cornified mouthparts; stomodeum vs. cornified mouthparts). Genes with an adjusted P-values (p-adj) < 0.05 and absolute log2 fold change (LFC) > 1 were considered significantly differentially expressed (DE), while all other genes were considered unbiased. This resulted in 2,123 unique DE genes. We identified DE genes of interest by searching for our HMM keratin hits, the AEDC genes reported in Sachslehner et al. (2026; their Table S1), and additional genes known to be involved in amniote hard cornification (e.g., SFTPs, EDC regulatory genes, transglutaminase and sulfhydryl oxidase enzyme genes, *Hoxc13*; Holthaus et al., 2025). We searched for the orthologs of *Xenopus SFTP1* and *SFTP2* in the *B. bombina* genome using a TBLASTN search.

### 2.7 Gene expression clustering, gene ontology and functional group enrichment analysis

For each stage of mouth development, we calculated the median normalized expression of the DE genes to include them in functional group enrichment analysis. We used MFuzz (v. 2.64.0; Kumar and Futschik 2007) to assign DE genes to expression clusters through time. For this, median normalized expression data were filtered for missing values and those with a standard deviation of 0 using the “filter.NA” and “filter.std” function of MFuzz. All DE genes passed this filter. Genes were soft clustered based on expression levels across the three developmental stages. Gene expression values were standardized to have a mean value of zero and a standard deviation of one with Mfuzz function “standardise”. The mFuzz “mestimate” function was used to calculate the fuzzification value (m=3.68) for optimal clustering. Genes with a membership value >= 0.7 were assigned to the same cluster. After visual inspection of different cluster numbers, we ultimately produced six clusters to maximize the number of unique expression profiles and collapse clusters with redundant expression patterns.

Gene ontology (GO) terms and GO categories were obtained using the R packages GO.db (v. 3.19.1; Carlson et al. 2019) and AnnotationDbi (v. 1.66.0; Pagès et al. 2024) to create a GO annotated database based on the reference genome annotation (GCF_027579735.1, BioProject: PRJNA915412). GO term enrichment analysis of biological processes (BP), cellular component (CC), and molecular function (MF), was carried out for each cluster with the R package clusterProfiler (v4.12.2; Wu et al. 2021) with all expressed genes defined as background genes. Functional terms with a *P*-value <0.05 (FDR adjusted using the Benjamini and Hochberg method) were considered enriched.

### 2.8 Histology

We investigated keratin and AEDC gene expression patterns in the jaw sheaths and keratodonts of *Bombina variegata* via *in situ* hybridization. All *B. variegata* tadpoles obtained for this study were produced from a breeding colony maintained at the University of Dayton (IACUC protocol # 024-05). Tadpoles were euthanized using buffered MS-222 and fixed in 4% paraformaldehyde for 24 hours at 4°C. Following fixation, the tadpoles were dehydrated through a DEPC PBC to ethanol gradient and then stored in 100% ethanol at -20°C until being prepared for paraffin sectioning. Specimens were decalcified in a 10% EDTA solution for 48 hours, embedded in paraffin following standard protocols, and sectioned to 8 μm in sagittal planes using a rotary microtome at the University of Dayton (on a Leica RM2155). The resulting tissue sections were then processed for *in situ* hybridization assays to characterize gene expression patterns.

### 2.9 RNA probe synthesis

We designed and synthesized RNA probes targeting a *krt34* paralog encoding a type I hard keratin, a *krt59* paralog encoding a type II hard keratin, and the cysteine-enriched *aedc15*. Because many of the *krt34* paralogs, *krt59* paralogs, and AEDC genes displayed similar expression profiles in *B. orientalis*, we selected one differentially expressed paralog from each family as a representative for evaluating spatial expression patterns during oral disk formation. Candidate *Bombina variegata* loci were identified by searching the *B. variegata* draft genome (NCBI assembly GCA_964205295.1) for *B. bombina* protein sequences that correspond to DE genes using TBLASTN: XP_053554654.1 (LOC128645590) for a *krt34* paralog, XP_053564318.1 (LOC128654436) for a *krt59* paralog, and XP_053547601.1 for *aedc15* (LOC128639486). The best-matching genomic regions were used to design riboprobe templates, and candidate amplicons were checked by BLAST to assess specificity relative to nearby paralogs and other keratin genes in *B. variegata*. Riboprobe templates were amplified using *B. variegata* cDNA sourced from a Gosner stage 24 tadpole using the following primer pairs: *krt34* paralog (forward 5’-ACAATGTCTCAGACTCGTTCCC-3’; reverse 5’-TTGTGCTTGAAGCTCGATCTCT-3’), *krt59* paralog (forward 5’-TGACCCAAACATCCAGAGAGTG-3’; reverse 5’-GTTCCTCTGCCTCCCTTATAGC-3’), and *aedc15* (forward 5’-ATGTCTGGAGTCAAAGGAGGACA-3’; reverse 5’-ATCTTTTTTCTTTCCACAAGGG-3’). A T7 promotor sequence (5’-TAATACGACTCACTATAGGG-3’) was added to the reverse primer for antisense probe synthesis. Digoxigenin-labeled antisense RNA probes were synthesized by *in vitro* transcription of PCR templates with T7 RNA polymerase (Promega) and DIG RNA Labeling Mix (Roche).

### 2.10 Section in situ hybridization

*In situ* hybridizations were performed on the paraffin-embedded tissue sections using the following protocol: tissue section slides were deparaffinized with HistoClear II, rehydrated, and then incubated in a prehybridization buffer for 2 hours. The DIG-labeled antisense RNA probes were denatured at 95°C for 12 minutes immediately before being applied to the tissue sections, which were then incubated at 61°C overnight. The following day, the slides underwent washes in 2× SSC and 0.5× SSC at 61°C and then incubated for 1 hour in blocking solution (2% Roche Blocking Reagent in maleic acid buffer containing Tween 20 [MABT]) at room temperature. Slides were antibody-labeled overnight at 4°C with anti-DIG-ALP (0.5 μL/mL; Roche). Following six MABT washes, substrate color reactions were carried out by adding 300 µL of BM purple (Roche) onto each slide at room temperature and replenished every 24 hours until the reactions were sufficient to visualize gene expression but minimized background signal. Slides were then fixed for 1 minute in 4% PFA, washed in PBS and distilled water, and mounted with Fluoromount-G with DAPI (Thermo Fisher), and imaged using an Olympus BX53 LED compound light microscope and Olympus DP23 camera. *In situ* hybridizations were performed multiple times to ensure that expression patterns were reproducible.

## 3 Results

### 3.1 Characterization of keratin genes in *Bombina*

In total, our HMM search identified 67 keratin genes within the *Bombina bombina* genome assembly, represented by 43 type I and 24 type II keratin genes (Table S2). All type I keratins resided on chromosome 1, with the exception of *krt18* on chromosome 3 and LOC128643914 on an unplaced scaffold, and all type II keratins resided on chromosome 3, excluding LOC128645268 on chromosome 1. The vertebrate keratin type I phylogeny recovered 13 *B. bombina* keratins and 30 *L. leishanense* keratins in a clade containing the single *krt34* orthologs from *Xenopus* and the axolotl (Figure 2A). This amphibian *krt34* clade was strongly supported as sister to the type I amniote hard keratins (UFBoot = 100). Many of these *Bombina* and *Leptobrachium* hard keratin sequences (hereafter named *krt34* paralogs) formed species-specific subclades, consistent with lineage-specific duplication. However, several *Bombina*-specific subclades were more closely related to *Leptobrachium-*specific subclades than to other *Bombina krt34* paralogs. Our keratin type II phylogeny recovered 12 *B. bombina* keratins and 30 *L. leishanense* keratins in a clade containing the single *krt59* orthologs from *Xenopus* and the axolotl (Figure 3A), as well as the type II amniote hard keratins (UFBoot = 99). Many of the *Bombina* and *Leptobrachium* hard keratin sequences (hereafter named *krt59* paralogs) formed species-specific subclades, however, like the *krt34* paralogs, the topology suggests some hard keratin duplications predated the divergence of these two genera. All *krt34* paralogs were adjacent within the type I keratin cluster on chromosome 1 (Figure 2B), and all *krt59* paralogs are adjacent within the type II keratin cluster on chromosome 3 (Figure 3B). This genomic organization supports their identification as true *krt34* and *krt59* paralogs and suggests that each group expanded through tandem duplication. Unlike the hard keratins of amniotes, none of the *krt34* and *krt59* paralogs in *B. bombina* are predicted to be enriched for cysteine residues (0–2.03% cysteine; Table S2).

**FIGURE 2.**
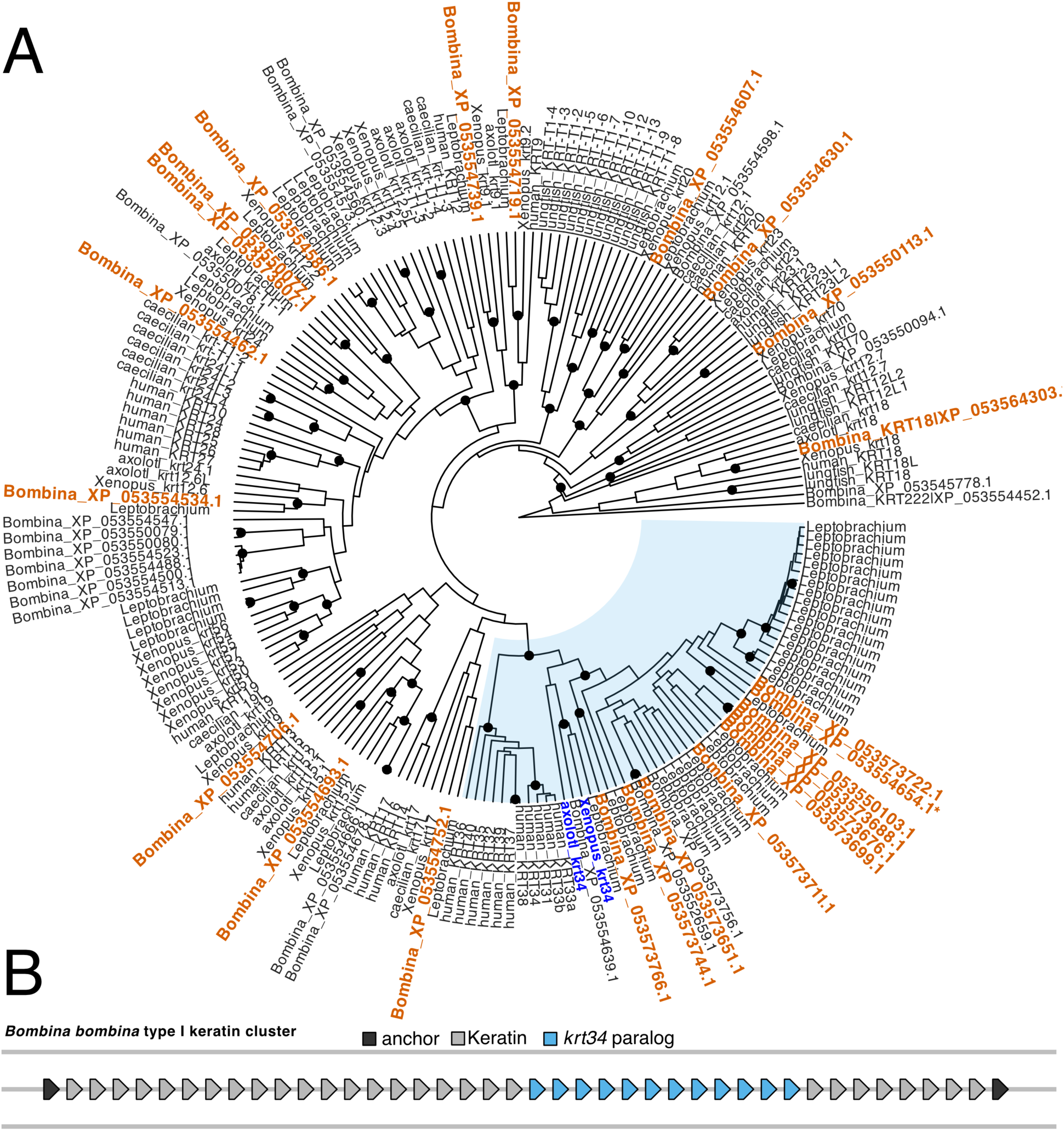
(A) Phylogeny of type I keratins. The clade highlighted in blue contains the tetrapod type I hard keratin homologs. *Bombina bombina* keratins highlighted in orange text are upregulated during oral disc formation. The *krt34* orthologs in *Xenopus tropicalis* and axolotl (*Ambystoma mexicanum*) are highlighted in blue text. Black nodes indicate bootstrap support ≥95. The asterisk indicates the *krt34* paralog corresponding to the RNA probe. *Leptobrachium leishanense* sequence names are available in Figure S4. (B) Spatial organization of type I keratin genes in *B. bombina* on chromosome 1. Gene order is preserved but sizes and spacing are not to scale. Adjacent *krt34* paralogs are denoted in blue.

**FIGURE 3.**
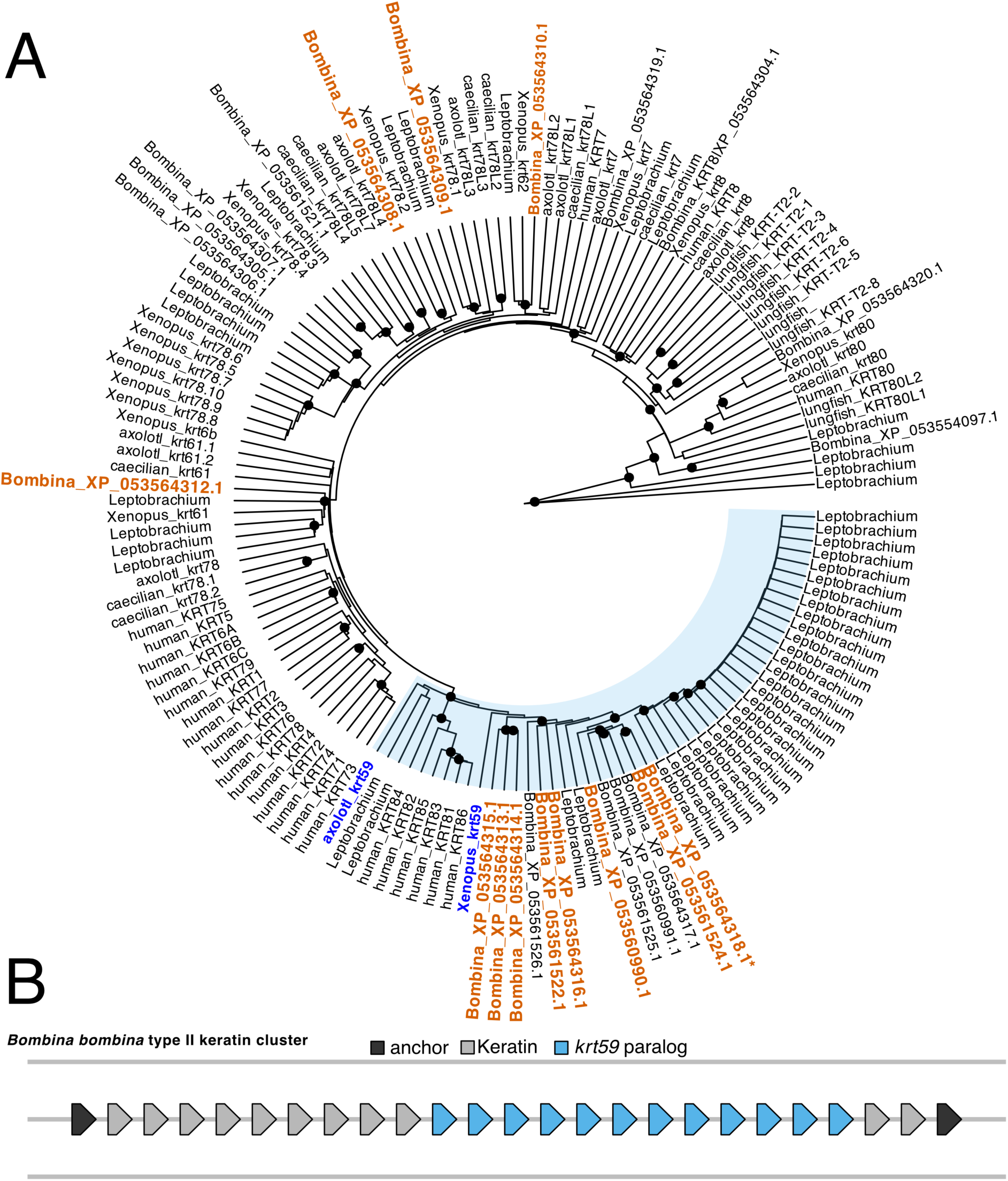
(A) Phylogeny of type II keratins. The clade highlighted in blue contains the tetrapod type II hard keratin homologs. *Bombina bombina* keratins highlighted in orange text are upregulated during oral disc formation. The *krt59* orthologs in *Xenopus tropicalis* and axolotl (*Ambystoma mexicanum*) are highlighted in blue text. Black nodes indicate bootstrap support ≥95. The asterisk indicates the *krt59* paralog corresponding to the RNA probe. *Leptobrachium leishanense* sequence names are available in Figure S5. (B) Spatial organization of type II keratin genes in *B. bombina* on chromosome 3. Gene order is preserved but sizes and spacing are not to scale. Adjacent *krt59* paralogs are denoted in blue.

### 3.2 Transcriptome profiles of oral disc development

Global gene expression differentiated the three developmental stages of oral disc formation (stomodeum, during oral disc formation, and cornified mouthparts) in *B. orientalis*, and each stage formed developmental-specific clusters in a principal component analysis (PCA; Figure S2). Gene expression differed more strongly between the stomodeum and oral disc formation (1,855 DE genes; Table S3) than between oral disc formation and fully cornified mouthparts (145 DE genes; Table S4). Differential gene expression was mostly driven by upregulation during oral disc formation in both comparisons (1,417 genes in the first comparison, 85 in later).

We identified 36 DE keratin genes, of which 33 (22 type I; 11 type II) were DE in the comparison between the stomodeum versus oral disc formation, and 13 (9 type I; 4 type II) were DE in the comparison between oral disc formation versus cornified mouthparts. Most of the DE keratins between the two comparisons represent the same genes expressed in the same direction, increasing across the three development time points (Figure 4). Ten of the 13 *B. bombina krt34* paralogs were DE in our RNAseq dataset. Of the 22 DE type I keratins in the comparison between stomodeum versus oral disc formation, two were upregulated in the *B. bombina* stomodeum (orthologs of *Xenopus krt70* and *krt18*) and 20 were upregulated during oral disc formation, including 10 *krt34* paralogs. Of the nine DE type I keratins in the comparison between oral disc formation versus cornified mouthparts, two were upregulated during oral disc development, and seven were upregulated in cornified mouthparts, including five *krt34* paralogs. The DE type I keratins outside of the *krt34* clade were widely distributed across the phylogeny and are sister to characterized *Xenopus* keratins (Table S2). Eight of the 12 *B. bombina krt59* paralogs were DE in our RNAseq dataset. All 11 of the DE type II keratins in the comparison between stomodeum versus oral disc formation were upregulated during oral disc formation, including eight *krt59* paralogs. All four of the DE type II keratins in the comparison between oral disc formation versus cornified mouthparts were upregulated in the cornified mouthparts, including three *krt59* paralogs. The four DE type II keratins outside of the *krt59* clade were orthologs to *Xenopus krt61, krt62*, *krt78.1*, and *krt78.2* (Table S2).

**FIGURE 4.**
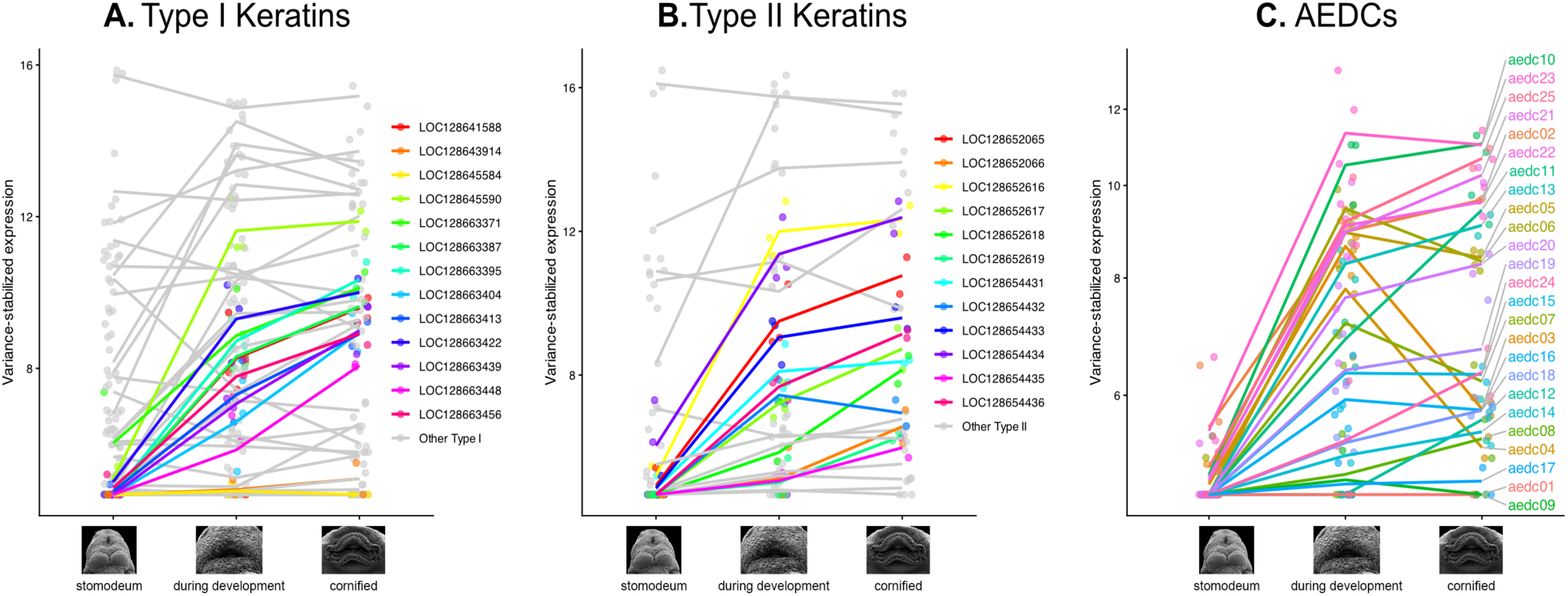
Expression profiles of (A) type I keratin genes, (B) type II keratin genes, and (C) AEDC genes across three developmental time points using variance-stabilized transformed counts (VST). Points represent individual biological replicates and lines connect the mean VST expression of each gene during stomodeum formation (developmental stage A), during oral disc formation (developmental stage B), and after mouthpart cornification (developmental stage C). Colored lines in (A) and (B) represent *krt34* and *krt59* paralogs, respectively. The *krt34* and *krt59* RNA probes targeted LOC128645590 and LOC128654436.

### 3.3 Identification of DE AEDCs and other cornification genes

In the comparison between the stomodeum versus oral disc formation, 17 AEDC genes (*aedc2*, *aedc3*, *aedc4*, *aedc5*, *aedc6*, *aedc7*, *aedc10*, *aedc11*, *aedc13*, *aedc15*, *aedc16*, *aedc19*, *aedc20*, *aedc21*, *aedc22*, *aedc23*) were upregulated during oral disc formation (Figure 4). In the later comparison, two AEDCs were upregulated during oral disc development (*aedc3*, *aedc4*) and two AEDCs were upregulated in cornified mouthparts (*aedc11*, *aedc25*). In addition to keratins and AEDCs, we identified *B. bombina* LOC128650459 and LOC128642747 as the orthologs of *Xenopus sftp1* and *sftp2*, respectively, which were upregulated in the oral disc compared to the stomodeum. At least two known EDC regulatory genes, *grhl3* and *klf4*, are also upregulated in tissues undergoing oral disc formation. *qsox1*, which encodes Quiescin sulfhydryl oxidase enzyme 1 that catalyzes disulfide bonds, is not DE among either of our comparisons but is expressed in all samples. *tgm1*, which encodes transglutaminase enzyme 1 that catalyzes isopeptide bonds, was upregulated during oral disc formation. *Hoxc13* was not DE among either of our comparisons but expressed in all samples. However, *foxn1*, a direct transcriptional downstream target of *Hoxc13* during hail and nail differentiation (Potter et al. 2011), was upregulated in samples undergoing oral disc formation compared to stomodeum samples.

### 3.4 Gene cluster expression dynamics

Among six MFuzz clusters of developmental gene expression changes, Cluster Two (283 genes) and Cluster Four (364 genes) captured expression profiles with expression increases in the same direction through development (Figure S3), either gradually (Cluster Two) or with a rapid increase between the stomodeum and oral disc formation (Cluster Four). Our GO analyses supported that Cluster Two was composed of genes involved in cell adhesion, keratinization, intermediate filament organization, hydrogen peroxide catabolic process, muscle contraction, oxygen transport, peptide cross-linking, calcium-dependent cell-cell adhesion via plasma membrane, cell adhesion molecules adherens junction organization, and regulation of membrane potential. Cluster 4 was characterized by genes involved in the process of keratinization and intermediate filament organization. The remaining clusters were associated with other biological processes (Table S5). Although keratin genes overall were present in all clusters except Cluster Three, all *krt34* and *krt59* paralogs were within Clusters Two and Four (Table S2). Cluster Two included nine *krt34* paralogs, six *krt59* paralogs, and six AEDCs (*aedc11*, *aedc13*, *aedc19*, *aedc21*, *aedc24*, *aedc25*), whereas Cluster Four included two *krt34* paralogs, four *krt59* paralogs, and seven AEDCs (*aedc2*, aedc5, *aedc10*, *aedc15*, *aedc16*, *aedc20*, *aedc22*). Four additional type I keratins and one type II keratin were in Cluster Two, and three additional type I keratins and two additional type II keratins were in Cluster Four (Table S2).

### 3.5 Histology and spatial gene expression patterns

The histology of the jaw sheath in *Bombina variegata* is comparable to that of other typical tadpoles, composed of epithelial columnar basal keratinocytes, stratified, flattened sheath cells, and stacked columns of cone-shaped cells (Figure 5A). Unlike most tadpoles, *Bombina* larvae can develop biserial keratodont rows, possessing double tooth rows within each labial ridge (Figure 5B). During the process of hard cornification, the apicalmost jaw sheath cells, keratodont cells, and body tubercle cells in *B. variegata* become black or dark brown in coloration (Figure 5A–C).

**FIGURE 5.**
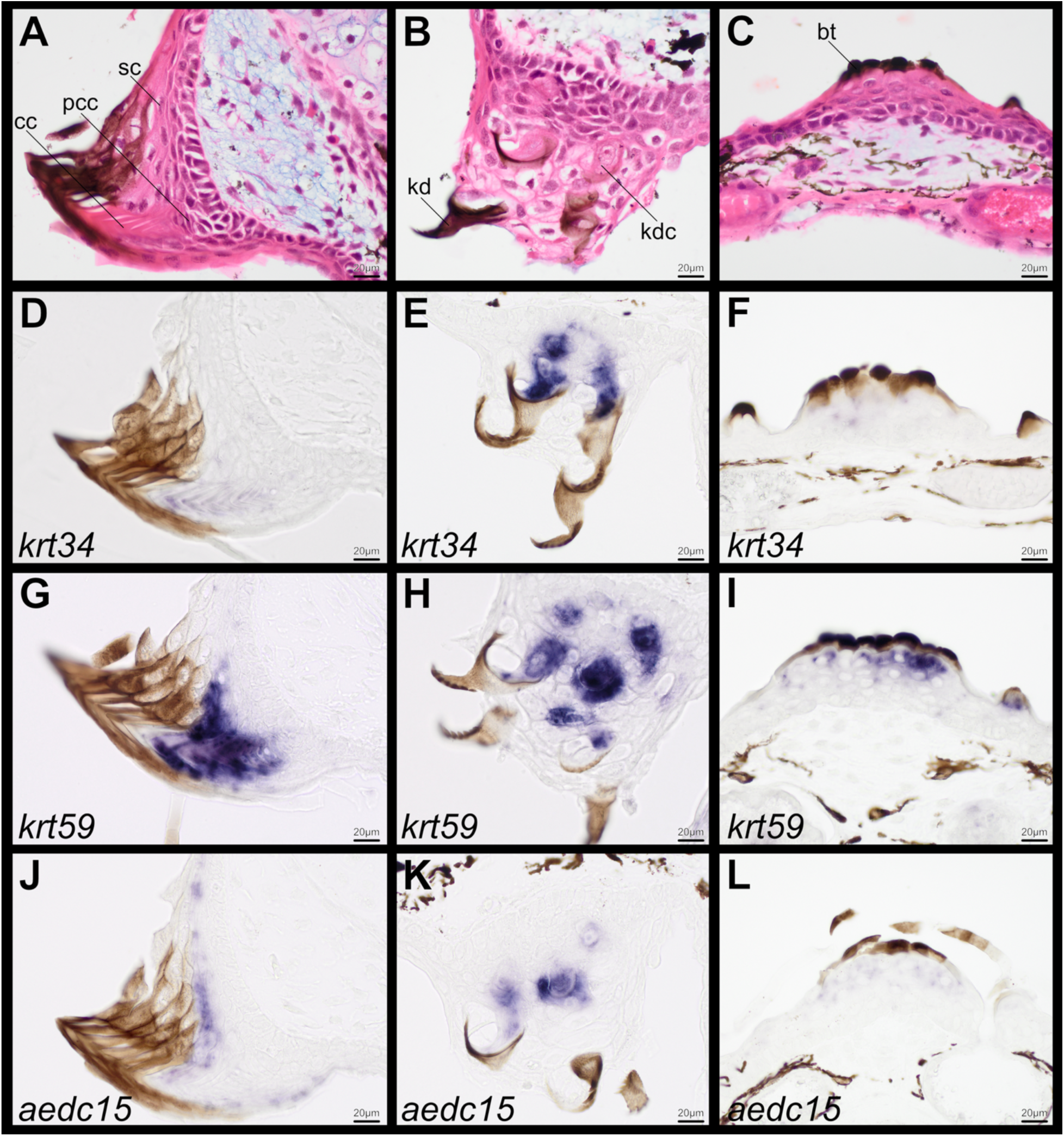
Sagittal sections through the tadpole cornified lower jaw sheath (left), biserial labial ridge with two keratodont loci (middle column), and froglet body tubercles (right) of *Bombina variegata* stained with H&E plus Alcian blue (A–C) and *in situ* hybridization (D–L). The *krt34* paralog (LOC128645590; D–F), *krt59* paralog (LOC128654436; G–I), and *aedc15* (J–L) are positively expressed in cells undergoing hard cornification. Abbreviations: bt, body tubercle; cc, cone cells; kd, keratodont; kdc, keratodont cell; pcc, precone cells; sc, sheath cells.

The *krt34* paralog (LOC128645590), the *krt59* paralog (LOC128654436), and *aedc15* were all expressed in the hard cornified mouthparts of *B. variegata* (Figure 5). The *krt34* paralog was weakly expressed in the central column of cone cells within the jaw sheaths (Figure 5D) yet was strongly expressed in multiple keratodont cell layers that are undergoing morphogenesis (Figure 5E). In comparison, the *krt59* paralog was strongly expressed in the central column of precone cells, cone cells, and adjacent sheath cells of the jaw sheaths (Figure 5G) and in multiple keratodont cell layers (Figure 5H). *aedc15* was expressed in both jaw sheath and keratodont cells (Figure 5J–K), but unlike *krt34* and *krt59*, *aedc15* had a peripheral expression pattern where only the apicalmost sheath and cone cell layers that are next in line to replace the terminalmost, darkened corneocytes are positive. The *krt59* paralog was strongly expressed in the body tubercle corneocytes of *B. variegata* froglets (Figure 5I), whereas *aedc15* and the *krt34* paralog were weakly expressed in these structures (Figure 5F, L). We did not detect expression for any of these genes in the soft cornified integument of *B. variegata* tadpoles or froglets.

## 4 Discussion

Our results demonstrate that, unlike *Xenopus*, both *Bombina* and *Leptobrachium* possess many duplicate copies of *krt34* and *krt59* paralogs that are orthologous to the type I and type II hard keratins of amniotes, respectively. Eighteen *B. bombina krt34* and *krt59* paralogs were upregulated in the oral disc during the formation and cornification of the jaw sheaths and keratodonts. Compared to *Bombina*, *Leptobrachium leishanense* has an expanded number of hard keratin genes, many of which are known to be expressed in the the prominent keratinized spines present in males along the upper jaw (Li et al. 2019). *Bombina* and *Leptobrachium* may have experienced some lineage-specific expansions of the *krt34* and *krt59* subclusters, although the topology of our type I and type II phylogenies indicated that several hard keratin duplications predated the divergence of *Bombina* and *Leptobrachium*, which last shared a common ancestor over 150 mya (Feng et al. 2017).

*Xenopus* could thus be unusual among frogs for possessing single copies of *krt34* and *krt59* and may have evolutionarily lost many duplicate hard keratin copies. All pipoid frogs, including *Xenopus*, secondarily lost cornified mouthparts during the tadpole phase (Figure 1B; Haas 2003), which may correspond to the reduction of *krt34* and *krt59* copies to a single gene each. *Xenopus* deploys *krt34* and *krt59* during hard cornification of claws (Carron et al. 2024) but only *krt59* was detected in adult nuptial pads (Steinbinder et al. 2026). These results indicate that two hard keratins may be sufficient for cornification of *Xenopus* claws, a single hard keratin may be sufficient for cornification of *Xenopus* nuptial pads, but many hard keratin copies may be necessary for the hard cornification of tadpole mouthparts. At least a few orthologs of amniote hard keratins are also expressed in frog toe pads, including one type I and two type II hard keratins in the toe pads of *Hyla cinerea* (Vandebergh et al. 2013) and one type I hard keratin in the toe pads of *Rhacophorus kio* (Wu et al. 2022). Toe pad cornification remains poorly understood in frogs, although Carron et al. (2024) proposed that the ancestral function of hard keratins in tetrapods may have been associated with hardening of the toe tips. The expanded keratin repertoire of amphibians relative to fishes, documented primarily in frogs, has been attributed to adaptations associated with the water-to-land transition of tetrapods (Vandebergh and Bossuyt 2012; Alibardi 2016; Ho et al. 2022). Our results instead suggest that this expansion was also driven, at least in part, by the ancient origin of the novel keratinized feeding apparatus of aquatic tadpoles (Chuliver et al. 2024).

The individual role of each *krt34* and *krt59* paralog in *B. bombina* tadpole mouthparts remains to be investigated. However, because many are expressed simultaneously and with the same developmental trajectory during oral disc development (Figure 4), a potential functional consequence of these duplications could be an increased capacity for keratin production. Higher rates of keratin synthesis could facilitate the rapid growth and continual replacement of jaw sheath and keratodont cells, which experience substantial mechanical wear during feeding and high cellular turnover rates (Kaung and Kollros 1977; Alibardi 2025). These duplications could also increase the density or complexity of the filament scaffold, impacting the material properties of keratinized mouthparts, such as increasing hardness, stiffness, or tensile strength. In addition to pipoids, there are several other anuran clades that have convergently lost larval jaw sheaths and keratodonts, including most microhylids (Haas 2003), several lineages of direct-developing frogs (Callery et al. 2001; Vera Candioti et al. 2025), and members of some specialized ecomorphological guilds, such as the sand-eating tadpoles (Dias et al. 2024, 2026). Their keratin gene repertoire should be characterized in a comparative context to further test this hypothesis. The axolotl (*Ambysoma mexicanum*) is similar to *Xenopus* in that it has only one *krt34* and *krt59*, but this neotenic salamander completely lacks hard cornified traits, which are generally rarely observed in salamanders. Hard cornified jaw sheaths are present in the genera *Pseudobranchus* and *Siren* (Sirenidae; Schwarz et al. 2021), hard cornified claws are restricted to the genus *Onychodactylus* ( Hynobiidae), and hard cornified body tubercles and nuptial pads are restricted to some members of the family Salamandridae (Holthaus et al. 2025).

Our expression results from one *krt34* and one *krt59* paralog show localized expression patterns associated with jaw sheath cells, keratodont cells, and body tubercle cells undergoing hard cornification in *B. variegata*. Neither gene was detected in the general integument of tadpoles or froglets. Both hard keratin genes are similarly expressed in keratodonts, whereas *krt59* was more strongly expressed in the jaw sheaths and body tubercles compared to *krt34*. Although jaw sheaths and keratodonts are both features of the oral disc, they differ substantially in morphology, function, and cellular dynamics. The jaw sheaths are the primary food removal apparatus and are usually arch-shaped with serrations on the edge (Altig 2007a). The keratodonts develop on the labial ridges and are hook-like structures disposed in a series of rows, anchoring the oral disc into the substrate, but also acting as food scrapers. Although the most common arrangement is two rows in the upper lip and three rows in the lower lip (denoted as a labial tooth row formula of 2/3), these numbers can vary up to 17/21 (Altig and Johnston 1989; Vera Candioti et al. 2025). As suggested by Dias and Haas (under review), differences in morphology of the jaw sheaths can be combined with differences in keratodont morphology and row formula in several ways, as a modular system. The spatial expression of additional *krt34* and *krt59* paralogs need to be characterized within each subcluster to test for distinct functions. Examining the number and expression of keratin copies in species with jaw sheaths but lacking labial teeth (e.g., *Cardioglossa* spp.; Hirschfeld et al. 2012), as well as those with many keratodont rows (e.g., *Ascaphus*; Vera Candioti et al. 2025), could also shed light on the differential roles of keratin genes across structures. In addition to the adult jaw spines in *L. leishanense*, the larval stage of this species also has more keratodonts rows (6/5 formula) and enlarged jaw sheaths compared to *Bombina* (Grosjean 2001), which could also be associated with its expanded hard keratin repertoire.

All upregulated *B. bombina* keratins in the oral disc exhibit low cysteine content (0–3%), similar to previous reports in *Xenopus* (Ehrlich et al. 2020). Despite the deployment of expanded hard keratins, tadpole mouthparts may ultimately become hardened and rigid through a structural reinforcement strategy distinct from the cysteine-enriched keratins of amniotes (Ehrlich et al. 2020). Alibardi (2010) predicted that hard cornification of the tadpole jaw sheath may be due to the deposition of undescribed, sulfur-rich proteins comparable to mammal keratin-associated proteins (KAPs), and this hypothesis has been supported by the detection of protein-bound sulfur in cornified nuptial pads and larval mouthparts (Luna et al. 2018; Dreger et al. 2025). A newly described, amphibian-specific gene family, the amphibian epidermal differentiation cluster (AEDC; Sachslehner et al. 2026), may encode Alibardi’s (2010) sulfur-rich proteins. *Bombina bombina* has 25 AEDCs located on chromosome 9 (Sachslehner et al. 2026), and of these, we identified 17 *B. orientalis* AEDCs upregulated in developing oral disc tissues, with most displaying expression trajectories similar to *krt34* and *krt59* paralogs (Figure 4). At least seven of the upregulated AEDCs (*aedc4*, *aedc15*, *aedc16*, *aedc19*, *aedc20*, *aedc21*, *aedc22*) encode short, cysteine-enriched (>15% cysteine residues; Sachslehner et al. 2026), low-complexity proteins that may be functionally analogous to mammalian KAPs in enabling disulfide cross-linking (Hirai et al. 2024). An additional four DE AEDCs (*aedc7*, *aedc10*, *aedc11*, *aedc13*) are highly enriched for glycine (16–44%; Sachslehner et al. 2026). Similar to our results, Sachslehner et al. (2026) also found that 14 AEDCs were upregulated in the larval mouth of *Bufo bufo* compared to tadpole skin, although they did not report fine-scale spatial expression patterns. Our results demonstrate that *aedc15*, a cysteine-enriched and upregulated AEDC in the *B. orientalis* oral disc, is specifically expressed in jaw sheath cells, keratodont cells, and body tubercle cells undergoing hard cornification in *B. variegata* but is absent from the general integument. Within the jaw sheaths, *aedc15* has a distinct, peripheral expression pattern that is restricted to cell layers that are directly beneath the terminalmost, darkened corneocytes.

Most hard cornified traits in frogs and salamanders are typically dark in coloration, often described as dark brown or black (Maddin et al. 2009). The source of this coloration has long been debated (Altig 2007b). Although these structures are frequently described as pigmented, several lines of evidence indicate that their dark color is not solely the result of melanin deposition (Altig 2007b; Maddin et al. 2009; Luna et al. 2018). For example, albino tadpoles and frogs retain darkened jaw sheaths, keratodonts, nuptial pads, and claws despite lacking normal melanin pigmentation (e.g., García-Diaz et al. 2023). Instead, the dark coloration appears to be intrinsically associated with the hard cornification process and may reflect the deposition of sulfur-rich structural proteins, the formation of disulfide cross-links, or both (Luna et al. 2018). If so, tissue color may serve as a proxy for material properties such as hardness or stiffness. Luna et al. (2018) proposed that the dark coloration of frog nuptial pads may be associated with the deposition of sulfur-rich structural proteins over the keratin intermediate filament scaffold. Consistent with this hypothesis, we found localized expression of *aedc15* within the next layer to undergo darkening in *B. variegata* jaw sheath cells. Further work is needed to characterize the spatial expression of additional AEDC copies with differing amino acid compositions. Testing whether protein-bound sulfur and disulfide cross-links colocalize with the peripheral expression of sulfur-rich AEDCs would provide a direct assessment of their proposed role in hard cornification.

The developmental genetic program underlying tadpole mouthpart formation remains poorly understood. These structures are ectodermal appendages (Holthaus et al. 2025; Paluh et al. 2025), and conserved developmental signaling pathways may regulate the induction and differentiation of the specialized jaw sheath, cone, and keratodont cell types. Recently, we showed that several key regulators of mineralized tooth development (another type of oral ectodermal appendage; Dhouailly et al. 2019), including *pitx2*, *sox2*, *ctnnb1*, and *dlx2*, are also expressed in both primordial and mature jaw sheath and keratodont tissues in tadpoles (Paluh et al. 2025). Notably, the expression domains of these regulatory genes are spatially distinct from those of *krt34*, *krt59*, and *aedc15*, being restricted to the basal and suprabasal epithelial cells at the base of the jaw sheath and keratodonts. This spatial division suggests that regulatory factors associated with oral/aboral epithelial patterning and maintenance are distinct from the differentiated cells that accumulate keratins and AEDCs as they undergo keratinization and cornification. Collectively, these findings indicate that tadpole keratinized mouthpart development is governed by a gene regulatory network comprising at least three functional components: (1) an oral/aboral ectodermal regulatory module that maintains the epithelium and specifies jaw sheath and keratodont cell fates, (2) a keratin intermediate filament module that establishes the structural scaffold, and (3) and a structural reinforcement module, which likely includes the amphibian epidermal differentiation cluster, whose proteins may harden the keratin scaffold through cross-linking. Further work is needed to characterize the developmental genetic program underlying tadpole keratinized mouthparts. Unanswered questions include: How do the genetic and cellular mechanisms governing jaw sheath *versus* keratodont development differ from one another? How do these programs compare with those underlying hard cornified traits in adult amphibians (e.g., nuptial pads, body tubercles, claws) and in amniotes? How has the program underlying tadpole feeding apparatus development evolved across anuran lineages that have diversified into distinct larval ecomorphological guilds with specialized feeding apparatus morphologies?

## Supporting Information

Additional supporting information can be found online in the Supporting Information section.

## Acknowledgments

We thank the Leibniz Institut zur Analyse des Biodiversitätswandels for supporting this research through the Innovation Fund. Dimitrij Trofimov and Felicitas Gunter facilitated our access to the clean lab of the Institut of Plant Science and Microbiology of the Universität Hamburg. We thank Julián Faivovich and Karina Silva Brandão for the enriching discussions on the subject. This work was supported by the U.S. National Science Foundation (DBI-2109344 to DJP) and by the University of Dayton Department of Biology.

## Author Contributions

**Conceptualization:** PHSD; DJP; **Methodology:** PHSD; DJP; PL; AB; AH; KH; NS; JJ; AC; SH; **Formal Analysis:** KH; PHSD; DJP; SH; IC; **Writing – Original Draft:** PHSD; KH; DJP; **Writing – Review & Editing:** PHSD; KH; NS; JJ; AC; SH; PL; IC; FVC; OH; AH; AB; DJP; **Project Administration:** PHSD; DJP; **Resources:** OH; AB; PL; AH; DJP; **Funding Acquisition:** PHSD; DJP; **Supervision:** DJP; AB; AH.

## Data Availability Statement

Data and scripts for all analyses are available on GitHub at https://github.com/dpaluh/tadpole_cornification.

## Ethics Statement

All embryos and tadpoles were reared in captivity at the Friedrich-Schiller-University Jena, Germany following their approved animal welfare protocol and at the University of Dayton (IACUC protocol # 024-05).

## Supporting Tables and Figures

**Table S1.** Total number of input reads and mapped reads.

**Table S2.** HMM keratin search results in *Bombina bombina*. Clade membership, cysteine richness, differential expression results, and mFuzz Cluster results are also summarized.

**Table S3.** Differential expression results in the comparison between stomodeum samples (before) vs. during oral disc formation. Positive log2FoldChange indicates upregulation during oral disc formation.

**Table S4.** Differential expression results in the comparison between during oral disc formation vs. cornified mouthparts (after). Positive log2FoldChange indicates upregulation in cornified mouthparts.

**Table S5.** Gene ontology (GO) enrichment analysis results for genes assigned to Mfuzz expression clusters.

**Figure S1.** Developmental stages of *Bombina* embryos used in this study. We recognized three developmental phases: stomodeum (A), during development (B), and cornified mouthparts (C). Note the development of keratinized features, such as the jaw sheaths and keratodonts (arrows in B).

**Figure S2.** (A) Transcriptome profiles of oral disc development. The three developmental groups can be differentiated along the PC1 axis (explaining 58% of the variance) with the cornified mouthparts (after) and oral disc formation (during) being closer to one another than the stomodeum samples (before). Moreover, cornified mouthparts can be clearly differentiated in the PC2 axis, which explains 16% of the variance. (B) Heatmap of the top 500 most highly expressed genes across different stages of oral disc formation. Gene expression levels are color-coded, with higher expression indicated in orange, and lower levels in blue; hierarchical clustering was performed on both genes and samples, with a resulting dendrogram clustering replicates by stage of development.

**Figure S3.** Soft clustering of differentially expressed genes. The X-axis characterizes the stage of mouth development, and the y-axis shows standardized and normalized expression of genes. Membership values of each gene are represented by color gradient, where genes with the highest membership values are represented in violet at the center of each distribution, and genes with the lowest in yellow.

**Figure S4.** Phylogeny of type I keratins with *Leptobrachium leishanense* sequence identifications. Black nodes indicate bootstrap support ≥95.

**Figure S5.** Phylogeny of type II keratins with *Leptobrachium leishanense* sequence identifications. Black nodes indicate bootstrap support ≥95.

