## Supplemental Figures S1-5 for "The molecular basis of hard cornification in the keratinized jaw sheaths and keratodonts of anuran tadpoles"

**FIGURE S1**. Developmental stages of *Bombina* embryos used in this study. We recognized three developmental phases: stomodeum (A), during development (B), and cornified mouthparts (C). Note the development of keratinized features, such as the jaw sheaths and keratodonts (arrows in B).

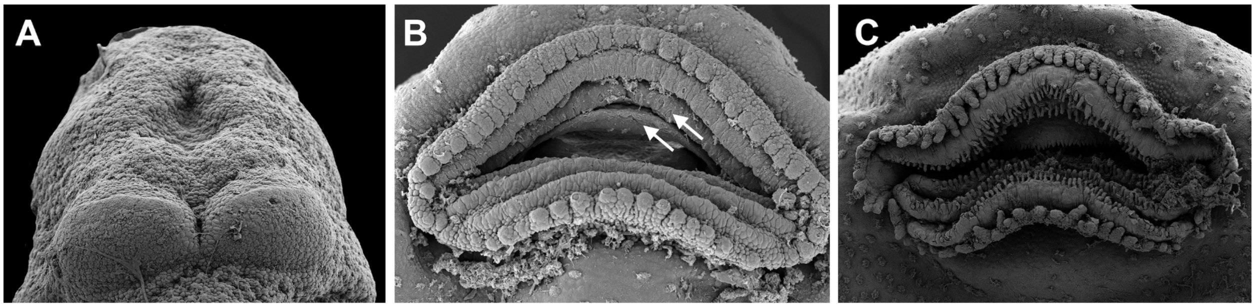

**FIGURE S2.** (A) Transcriptome profiles of oral disc development. The three developmental groups can be differentiated along the PC1 axis (explaining 58% of the variance) with the cornified mouthparts (after) and oral disc formation (during) being closer to one another than the stomodeum samples (before). Moreover, cornified mouthparts can be clearly differentiated in the PC2 axis, which explains 16% of the variance. (B) Heatmap of the top 500 most highly expressed genes across different stages of oral disc formation. Gene expression levels are color-coded, with higher expression indicated in orange, and lower levels in blue; hierarchical clustering was performed on both genes and samples, with a resulting dendrogram clustering replicates by stage of development.

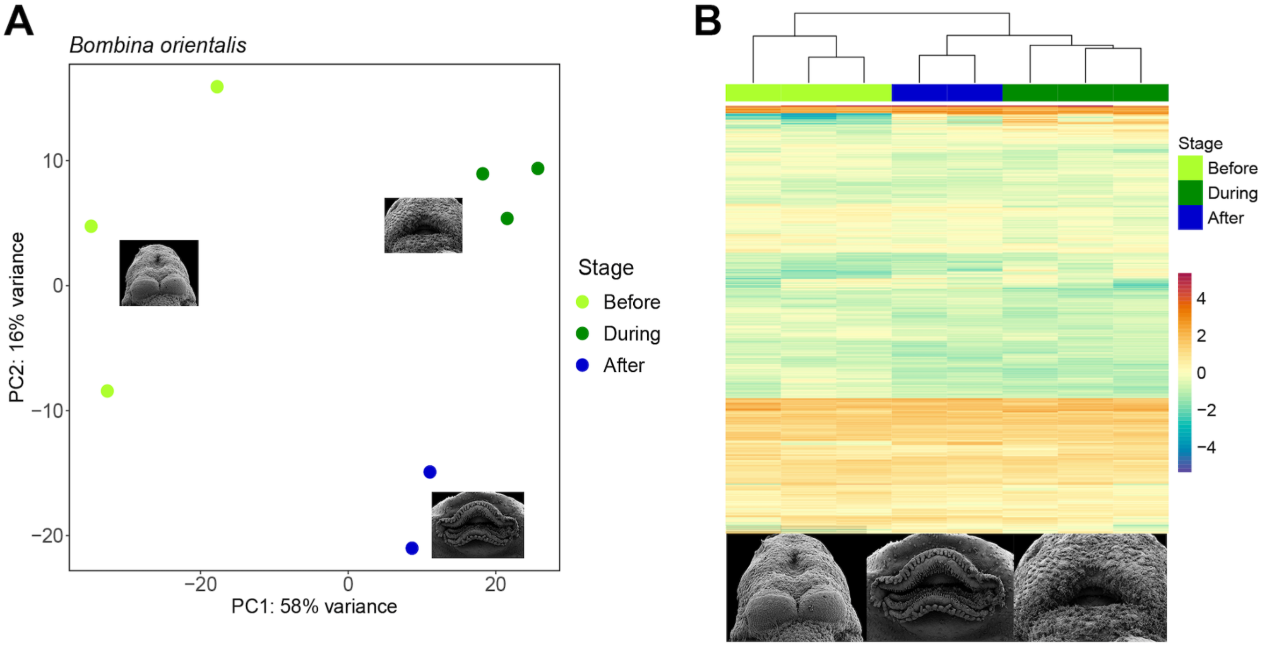

**FIGURE S3.** Soft clustering of differentially expressed genes. The X-axis characterizes the stage of mouth development, and the y-axis shows standardized and normalized expression of genes. Membership values of each gene are represented by color gradient, where genes with the highest membership values are represented in violet at the center of each distribution, and genes with the lowest in yellow.

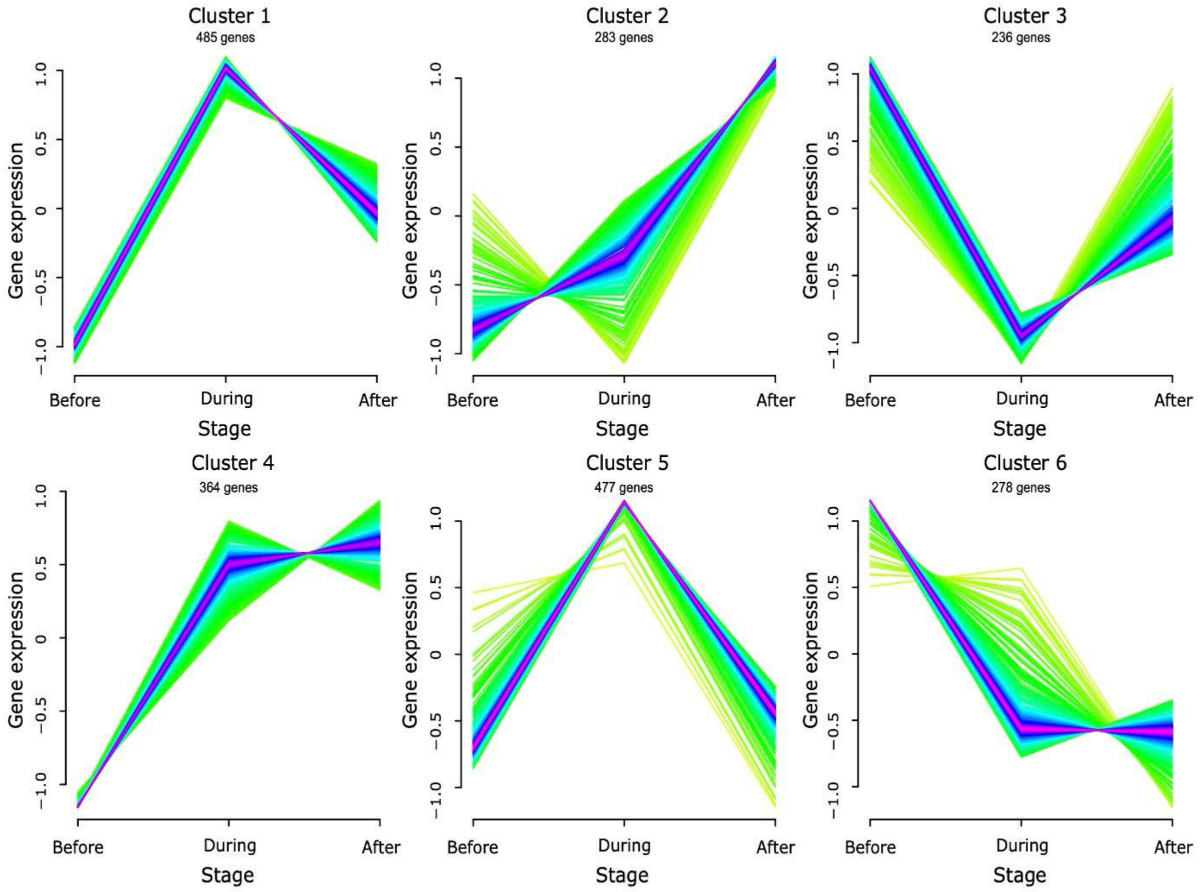

**FIGURE S4.** Phylogeny of type I keratins with *Leptobrachium leishanense* sequence identifications. Black nodes indicate bootstrap support ≥95.

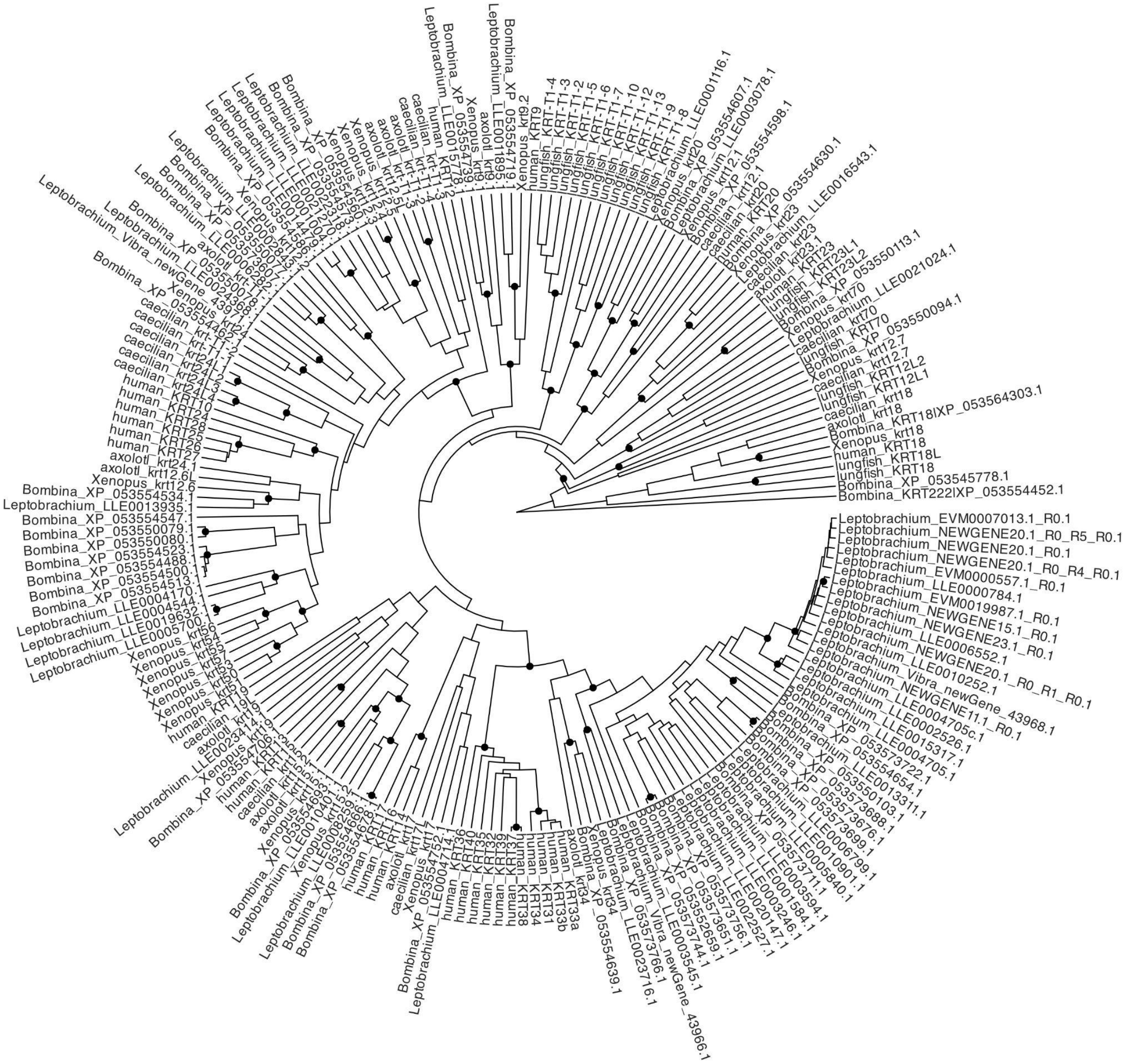

**FIGURE S5.** Phylogeny of type II keratins with *Leptobrachium leishanense* sequence identifications. Black nodes indicate bootstrap support ≥95.

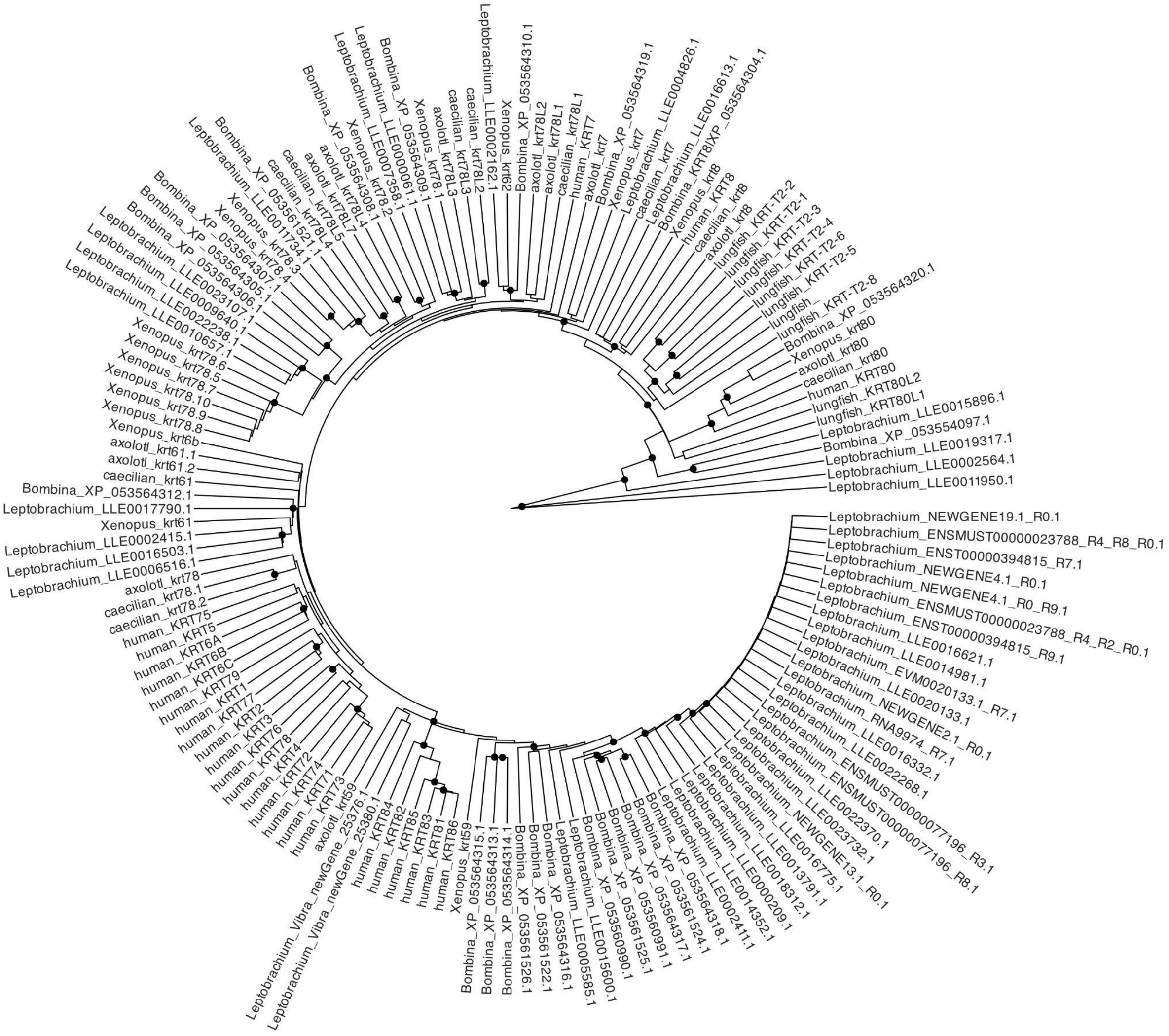
